# Bayesian genome-wide analysis of cattle traits using variants with functional and evolutionary significance

**DOI:** 10.1101/2021.05.05.442705

**Authors:** Ruidong Xiang, Ed J. Breen, Claire P. Prowse-Wilkins, Amanda J. Chamberlain, Michael E. Goddard

## Abstract

**Context:** Functional genomics studies have revealed genomic regions with regulatory and evolutionary significance. Such information independent of association analysis may benefit fine-mapping and genomic selection of economically important traits. However, systematic evaluation of the use of functional information in mapping, and genomic selection of cattle traits is lacking. Also, Single Nucleotide Polymorphisms (SNPs) from the high-density (HD) panel are known to tag informative variants, but the performance of genomic prediction using HD SNPs together with variants supported by different functional genomics is unknown.

**Aims:** We selected six sets of functionally important variants and modelled each set together with HD SNPs in Bayesian models to map and predict protein, fat, and milk yield as well as mastitis, somatic cell count and temperament of dairy cattle.

**Methods:** Two models were used: 1) BayesR which includes priors of four distribution of variant-effects, and 2) BayesRC which includes additional priors of different functional classes of variants. Bayesian models were trained in 3 breeds of 28,000 cows of Holstein, Jersey and Australian Red and predicted into 2,600 independent bulls.

**Key results:** Adding functionally important variants significantly increased the enrichment of genetic variance explained for mapped variants, suggesting improved genome-wide mapping precision. Such improvement was significantly higher when the same set of variants were modelled by BayesRC than by BayesR. Combining functional variant sets with HD SNPs improves genomic prediction accuracy in the majority of the cases and such improvement was more common and stronger for non-Holstein breeds and traits like mastitis, somatic cell count and temperament. In contrast, adding a large number of random sequence variants to HD SNPs reduces mapping precision and has a worse or similar prediction accuracy, compared to using HD SNPs alone to map or predict. While BayesRC tended to have better genomic prediction accuracy than BayesR, the overall difference in prediction accuracy between the two models was insignificant.

**Conclusions:** Our findings demonstrate the usefulness of functional data in genomic mapping and prediction.

**Implications:** We highlight the need for effective tools exploiting complex functional datasets to improve genomic prediction.

## Introduction

Emerging evidence shows that genomic variants with causal roles in biology can be used to improve genomic prediction of complex traits. The biological function of genomic variants provides information independent of genotype-trait associations which are usually confounded by linkage disequilibrium (LD). Such independent information can be exploited to identify informative variants. Once identified, informative variants can be used to improve genomic prediction (Xiang, MacLeod, Daetwyler, de Jong, O’Connor, Schrooten, Chamberlain & Goddard, 2021). While the use of functional data in improving genomic mapping and prediction has been reported in humans (Amariuta, Ishigaki, Sugishita, Ohta, Koido, Dey, Matsuda, Murakami, Price & Kawakami, 2020; Weissbrod, Hormozdiari, Benner, Cui, Ulirsch, Gazal, Schoech, Van De Geijn, Reshef & Márquez-Luna, 2020), using functional data in predicting the genetic merit of animal traits has not been comprehensively examined. However, there is evidence in cattle supporting the advantage of the use of functional information in genomic mapping and prediction with the linear mixed model (Fang, Sahana, Ma, Su, Yu, Zhang, Lund & Sørensen, 2017a; Fang, Sahana, Ma, Su, Yu, Zhang, Lund & Sørensen, 2017b; Liu, Fang, Zhou, Santos, Xiang, Daetwyler, Chamberlain, Cole, Li, Yu, Ma, Zhang & Liu, 2019; Xiang, Berg, MacLeod, Hayes, Prowse-Wilkins, Wang, Bolormaa, Liu, Rochfort, Reich, Mason, Vander Jagt, Daetwyler, Lund, Chamberlain & Goddard, 2019; Xu, Gao, Wang, Xu, Liu, Chen, Xu, Gao, Zhang & Gao, 2020).

The Functional Annotation of ANimal Genomes (FAANG) consortium (Clark, Archibald, Daetwyler, Groenen, Harrison, Houston, Kühn, Lien, Macqueen & Reecy, 2020) provides many types of sequencing data indicating the functionality of genome-wide sites (examples reviewed in (Clark *et al.*, 2020)). While these public datasets await exploitation, the structure and information content of different functional datasets vary significantly. For example, we recently showed that amongst all analysed functional datasets, a set of 300,000+ sequence variants within sites highly conserved across 100 vertebrate species had the strongest enrichment with cattle trait heritability (Xiang *et al.*, 2019), which primarily influences genomic prediction accuracy. Additionally, a few thousand variants affecting the concentration of milk fat metabolites, i.e., metabolic mQTLs, also had significantly higher variance than SNPs in the 50K panel for cattle traits. Millions of variants that change gene expression levels (geQTLs) or RNA splicing (sQTLs) are also enriched with complex trait QTL (Fink, Lopdell, Tiplady, Handley, Johnson, Spelman, Davis, Snell & Littlejohn, 2020; Li, van de Geijn, Raj, Knowles, Petti, Golan, Gilad & Pritchard, 2016; Lopdell, Tiplady, Struchalin, Johnson, Keehan, Sherlock, Couldrey, Davis, Snell & Spelman, 2017; Silva, Fonseca, Pinheiro, Magalhães, Muniz, Ferro, Baldi, Chardulo, Schnabel & Taylor, 2020; Xiang, Hayes, Vander Jagt, MacLeod, Khansefid, Bowman, Yuan, Prowse-Wilkins, Reich, Mason, Garner, Marett, Chen, Bolormaa, Daetwyler, Chamberlain & Goddard, 2018).

However, recent studies showed that variants close to genes with high or specific expression patterns had limited improvement in prediction accuracy (de Las Heras-Saldana, Lopez, Moghaddar, Park, Park, Chung, Lim, Lee, Shin & van der Werf, 2020; Fang, Cai, Liu, Canela-Xandri, Gao, Jiang, Rawlik, Li, Schroeder & Rosen, 2020). Another common type of functional data is peaks from ChIP-seq for histone modifications which are enriched with promoters and/or enhancers regulating gene activities (Carey, Peterson & Smale, 2009). Our work showed that hundreds of thousands of variants under ChIP-seq peaks are enriched for complex trait QTL in cattle (Prowse-Wilkins, Wang, Xiang, Goddard & Chamberlain, 2021; Xiang *et al.*, 2019). In addition, variants within the gene coding regions are expected to have a high impact on complex traits. However, we and others previously found coding-related variants (around 100,000) have limited contributions to cattle trait heritability (Koufariotis, Chen, Stothard & Hayes, 2018; Xiang *et al.*, 2019), although their use in improving genomic prediction has not been studied.

One way to assess the information content of functional data is to compare variants prioritised by functional data with SNPs from standard genotyping panels. We have previously performed such assessment using the standard 50K bovine SNP chip and showed that functional information can improve genomic prediction accuracy compared to the 50K chip SNPs (Xiang *et al.*, 2021). However, denser panels such as the high-density (HD) SNP chip containing ~700,000 SNPs across the genome may be able to tag many functional elements via LD, although it is not routinely used in animal genomic evaluation. With the development of animal breeding, the HD panel may be intensively used in the future genomic evaluation. Therefore, it is of interest to know if functional information can provide any advantage in genomic mapping and prediction when HD SNPs are used. Also, since causal variants are expected to have similar phenotypic effects across different breeds, we aim to compare the use of functionally important variants in genomic prediction across different breeds.

In the present study, we evaluate sequence variant sets prioritised by 6 types of functional and evolutionary data in combination with the standard HD SNPs in genomic mapping and prediction of 6 dairy cattle traits. We train the prediction equations using the BayesR method (Erbe, Hayes, Matukumalli, Goswami, Bowman, Reich, Mason & Goddard, 2012) which fits a mixture of 4 distributions of variant-effects and using the BayesRC method which fits different distributions for each functional class of variant classifications (MacLeod, Bowman, Vander Jagt, Haile-Mariam, Kemper, Chamberlain, Schrooten, Hayes & Goddard, 2016).

Genomic predictors were trained using 28,000 cows that included 3 breeds: Holstein, Jersey and Australian Red. Genomic estimated breeding values (gEBVs) were predicted and validated in 2,500 Holstein, Jersey and Australian Red bulls. We compare the results of mapping and genomic prediction across the above-described scenarios, discuss these results and provide suggestions for future studies.

## Materials and Methods

The phenotype data analysed in this study were collected by DataGene Australia (http://www.datagene.com.au/) and no further live animal experimentation was required for our analyses. A set of 28,049 Australian cows were used as the discovery population and a set of 2,567 bulls were used as the validation population. The bull phenotypes were obtained as daughter trait deviations: i.e. the average trait deviations of a bull’s daughters pre-corrected for known fixed effects by DataGene. The cow phenotypes were measured on themselves.

Note that these bulls and cows were not included in those 44,000+ animals used to discover functional variants (Xiang *et al.*, 2019; Xiang *et al.*, 2021; Xiang, van den Berg, MacLeod, Daetwyler & Goddard, 2020). We also checked the pedigree to make sure that bulls used in the validation population were not the sires of cows from the discovery population. Cows in the discovery set included 24,305 Holstein, 2,486 Jersey, 1,258 Australian Red. Bulls in the validation datasets contained 2,091 Holstein, 385 Jersey, 91 Australian Red. Traits considered in the analysis included protein yield (Prot), fat yield (Fat), milk yield (Milk), Mastitis (Mas), somatic cell count (Scc) and temperament (Temp).

The genotypes used in the study were imputed sequence variants based on Run7 of the 1000 Bull Genomes Project (Daetwyler, Capitan, Pausch, Stothard, Van Binsbergen, Brøndum, Liao, Djari, Rodriguez & Grohs, 2014; Hayes & Daetwyler, 2018) based on the ARS-UCD1.2 reference bovine genome (https://www.ncbi.nlm.nih.gov/assembly/GCF_002263795.1/) (Rosen, Bickhart, Schnabel, Koren, Elsik, Tseng, Rowan, Low, Zimin & Couldrey, 2020). Variants with Minimac3 (Fuchsberger, Abecasis & Hinds, 2014; Howie, Fuchsberger, Stephens, Marchini & Abecasis, 2012) imputation accuracy *R*^2^ > 0.4 and minor allele frequency (MAF) > 0.005 in bulls and cows. Most bulls were genotyped with a medium-density SNP array (50K) or a high-density SNP array and most cows were genotyped with a low-density panel of approximately 6,900 SNPs overlapping with the standard-50K panel (BovineSNP50 beadchip, Ilumina Inc). The low-density genotypes were first imputed to the Standard-50K panel and then all 50K genotypes were imputed to the HD panel using Fimpute v3 (Sargolzaei, Chesnais & Schenkel, 2014; Xiang *et al.*, 2019). Then, all HD genotypes were imputed to sequence using Minimac3 with Eagle (v2) to pre-phase genotypes (Howie *et al.*, 2012; Loh, Danecek, Palamara, Fuchsberger, Reshef, Finucane, Schoenherr, Forer, McCarthy & Abecasis, 2016). We aimed to test whether variant sets selected from different functional and/or evolutionary information, in addition to the standard HD SNP panel, can be useful for genomic prediction. Therefore, we first included a baseline set, which is 610,764 SNPs from the standard bovine high-density panel. There were six functional and/or evolutionary variant sets: 549,007 variants under multiple ChIP-seq peaks (Kern, Wang, Xu, Pan, Halstead, Chanthavixay, Saelao, Waters, Xiang & Chamberlain, 2021; Prowse-Wilkins *et al.*, 2021) (‘ChiPseq’), 106,538 variants annotated as related to coding activities by Ensembl Variant Effect Predictor (McLaren, Gil, Hunt, Riat, Ritchie, Thormann, Flicek & Cunningham, 2016) (‘Coding’), 943,315 variants affecting RNA splicing sQTLs from 4 cattle tissues (Chamberlain, Hayes, Xiang, Vander Jagt, Reich, Macleod, Prowse-Wilkins, Mason, Daetwyler & Goddard, 2018; Daetwyler, Xiang, Yuan, Bolormaa, Vander Jagt, Hayes, van der Werf, Pryce, Chamberlain & Macleod, 2019; Xiang *et al.*, 2018) (‘sQTL’), 65,394 finely mapped variants with pleiotropic effects genome-wide (Xiang *et al.*, 2021) (‘Finemap80k’), 4,871 variants affecting milk fat metabolites mQTLs (Xiang *et al.*, 2019) (‘mQTL’) and 317,279 conserved sites across 100 vertebrates (Xiang *et al.*, 2019) (‘Cons100w’). Note that some of these functional variant sets were initially determined on the UMD3.1 genome and were from different cattle populations. These sets were lifted over from the older genome to ARS-UCD1.2 and filtered with imputation accuracy and MAF in the new cattle populations.

The model training of the above-described data used BayesR (Erbe *et al.*, 2012) and BayesRC (MacLeod *et al.*, 2016), which are now implemented via BayesR3, with improved efficiency using blocks. BayesR jointly models all variants together with different effect distribution priors. BayesRC follows the same approach but in addition allows a ‘C’ prior which models classes of variants. Another aim is to see whether there are differences in genomic prediction accuracy by modelling the same variants using BayesR and BayesRC. To aid this comparison, we combined each functional variant set with the HD variants which led to 6 combined variant sets: 1) ChIP-seq peak tagged variants + HD SNPs (‘ChiPseq_HD’), 2) coding variants + HD variants (‘Coding_HD’), 3) sQTL variants + HD SNPs (‘sQTL_HD’), 4) finely mapped variants + HD SNPs (‘Finemap80k_HD’), 5) mQTL variants + HD SNPs (‘mQTL_HD’) and 6) conserved variants + HD SNPs (‘Cons100w_HD’). The average minor allele frequency of these sets of variants were 0.22 (±0.00014) for ChiPseq_HD, 0.25(±0.0002) for Coding_HD, 0.24 (±0.0001) for sQTL_HD, 0.27 (±0.0002) for Finemap80k_HD, 0.27 (±0.0002) for mQTL_HD, 0.23 (±0.0002) for Cons100w_HD, and 0.27 (±0.0002) for HD alone.

In single-trait BayesR, we directly model these 6 variant sets one set at a time. To create a reference baseline, we also used single-trait BayesR to fit the HD variant set (‘HD’) alone. In single-trait BayesRC, for each of the same 6 combined variant sets, we specify 2 different variant classes: 1) Variants appeared in the functional and/or evolutionary set and 2) variants only appeared in the HD variant set.

Both BayesR and BayesRC modelled variant effects as a mixture distribution of four normal distributions including a null distribution, *N*(0, 0.0σ^2^_*g*_), and three others: *N*(0, 0.0001σ^2^_*g*_), *N*(0, 0.001σ^2^_*g*_), *N*(0, 0.01σ^2^_*g*_), where σ^2^_*g*_ was the additive genetic variance for the trait. The starting value of σ^2^_*g*_ for each trait was estimated using GREML implemented in the MTG2 (Lee & Van der Werf, 2016) with a single genomic relationship matrix made of all sequence variants. The statistical model used in the single-trait BayesR and BayesRC in was:

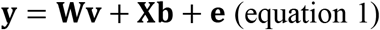

where **y** was a vector of phenotypic records; **W** was the design matrix of marker genotypes; centred and standardised to have a unit variance; **v** was the vector of variant effects, distributed as a mixture of the four distributions as described above; **X** was the design matrix allocating phenotypes to fixed effects; **b** was the vector of fixed effects, including breeds; **e** = vector of residual errors. As a result, the effect *b* for each variant jointly estimated with other variants were obtained for further analysis.

BayesRC used the same linear model as BayesR. The C component of BayesRC had two categories *c*(*c* = 2) as described above. Within each category *c*, an uninformative Dirichlet prior (α) was used for the proportion of effects in each of the four normal distributions of variant effects: *P*_*c*_~*Dir*(α_*c*_), where *a*_*c*_ = [1, 1, 1, 1]. α_*c*_ was updated each iteration within each category: *P*_*c*_~*Dir*(α_*c*_ + β_*c*_), where β_*c*_ was the current number of variants in each of the four distributions within category c, as estimated from the data.

Two metrics were evaluated for mapping results. One is the mixing proportion, i.e., the proportion of variants with small effect *N*(0, 0.0001σ^2^_*g*_), medium effect *N*(0, 0.001σ^2^_*g*_) and large effect *N*(0, 0.01σ^2^_*g*_) for each BayesRC run across the functional variant class and the HD SNP class. This metric shows the information content of the two classes. The other metric was the percentage of 50kb segments needed by the model to explain 50% of the cumulative sum of posterior probability (PP), which indicated the mapping precision. For each variant, PP was calculated as 1 – *P_0_* where *P_0_* was the probability for the variant to be within the zero-effect distribution *N*(0, 0.0σ^2^). The sum of PP across all variants estimates the number of variants causing genetic variance in the trait. The smaller amount of genomic segments needed to explain a cumulative sum of PP, the higher the mapping precision. We also compared genomic prediction accuracy, defined as the Pearson correlation *r* between genomic estimated breeding value (gEBV) and phenotype in the validation populations. gEBV of the validation animals was calculated as *gEBV* = ***Zŝ*** (equation 2), where ***Z*** was a matrix of the standardisd genotypes of animals in the validation set, and ***ŝ*** was the vector of variant effects from the training model. In addition, to test if adding a large number of random variants to the HD panel can increase mapping precision and prediction accuracy, a random set of 944,616 variants matching the size of the largest set of functional variants (sQTL, 943,315 variants) was also selected and added to the HD panel (‘Random_HD’). This random set was analysed for BayesR, mapping precision and prediction accuracy in the same fashion as other variant sets described above.

## Results

### Information content in the functional variant sets

Averaged across mixing proportions from single-trait BayesRC, we show that compared to HD SNPs, the finely mapped variants had consistently higher enrichment with variants showing small, medium and large effects (Figure 1). Variants within coding regions showed higher enrichment than HD SNPs for large- and medium-effect variants. Interestingly, mQTLs, which were variants affecting the concentration of milk fat metabolites (Benedet, Ho, Xiang, Bolormaa, De Marchi, Goddard & Pryce, 2019; Xiang *et al.*, 2019), had lower enrichment of small-effect variants than HD SNPs, but had higher enrichment of medium and large-effect variants than HD SNPs.

**Figure 1.**
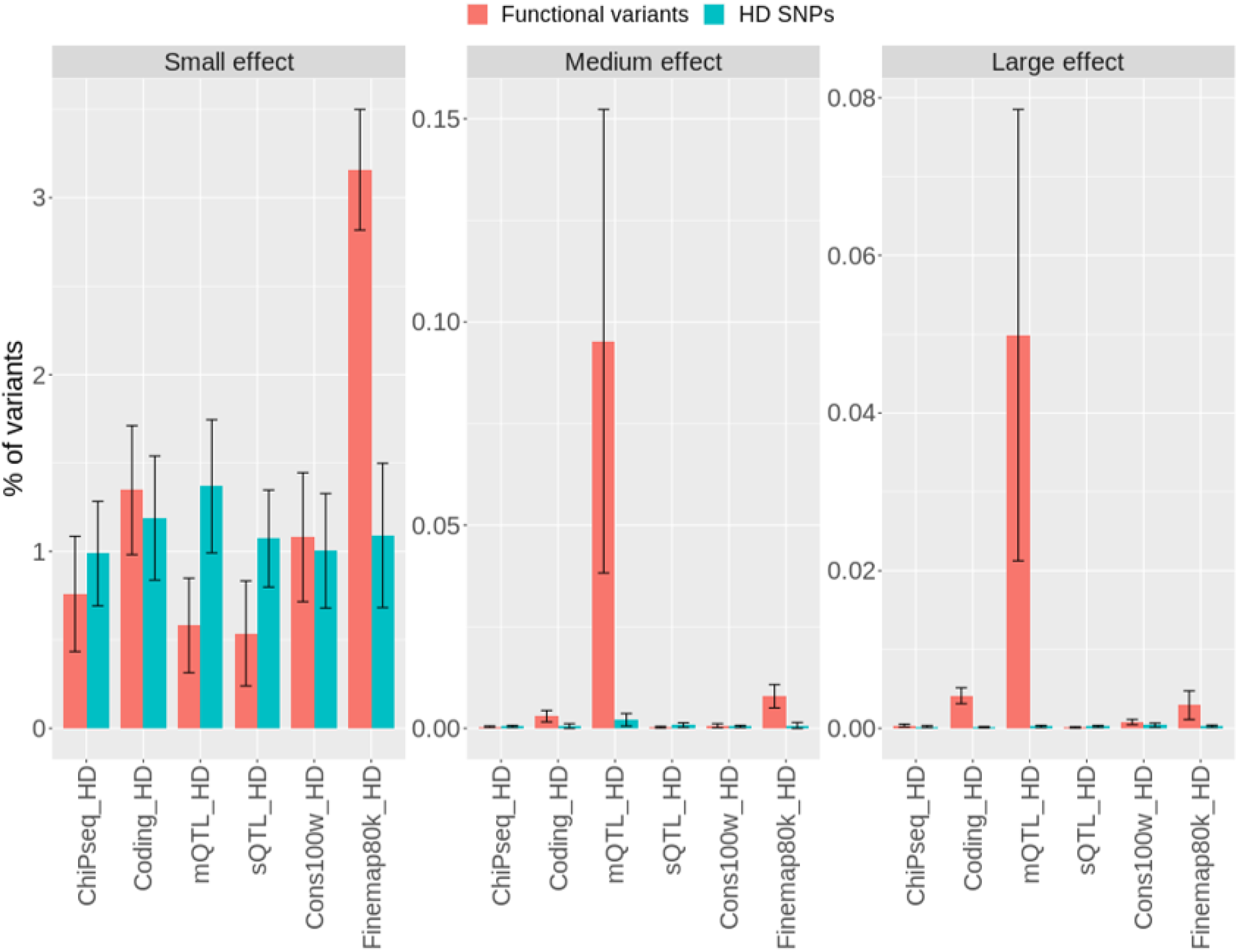
The proportion of small-effect, medium effect and large effect variants in functional variants and HD SNPs. The mean and standard error bars are averaged across 6 traits. ChiPseq_HD: ChIP-seq peaks + HD SNPs. Coding_HD: coding variants + HD SNPs. mQTL_HD: mQTLs + HD SNPs. sQTL_HD: sQTL variants + HD SNPs. Cons100w_HD: conserved variants across 100 vertebrates + HD SNPs. Finemap80k_HD: finely mapped variants + HD SNPs.

### Mapping precision

Across traits, we show that all models using functional variants, except mQTL, needed a smaller amount of genome-wide segments to explain 50% of the cumulative sum of PP, compared to HD SNPs (Figure 2). This means that when adding to the HD SNPs, most functional variants increased mapping precision. In contrast, adding randomly selected 944,000 variants to HD SNPs increased the amount of genome-wide segments (by 2.82%± 0.13%) across scenarios to explain 50% of the cumulative sum of PP, compared to only using HD SNPs. This suggested that adding random variants to HD decreases mapping precision. It is worth noting that when using 106,538 coding variants and 65,394 finely mapped variants, BayesRC provided a further increase in mapping precision over HD SNPs than BayesR. On the other hand, when using 549,007 ChIP-seq tagged variants and 943,315 sQTL variants, BayesRC had less increase in mapping precision over HD SNPs than BayesR. This could be due to the reduced signal-to-noise ratio in large variant sets of ChIP-seq tagged variants and sQTLs.

**Figure 2.**
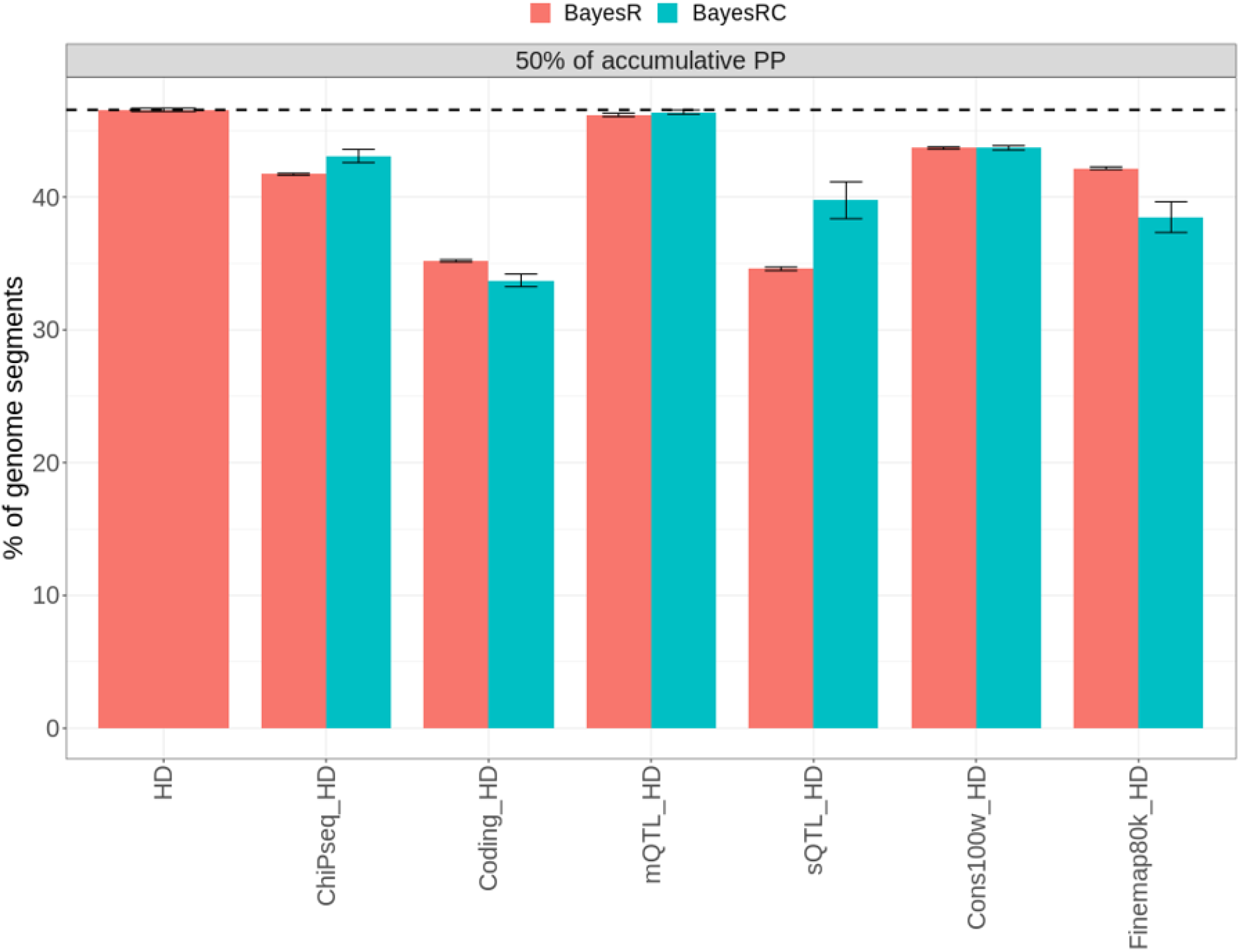
Mapping precision of different models. The Y-axes represent the percentage of 50kb segments needed by the model to explain 50% of the cumulative sum of posterior probability (PP) of variants. A shorter bar means less amount of segments the model needs to explain the same amount of genetic variance, indicating higher mapping precision. Black dashed line indicates the Y value for the HD SNPs, fitted along in BayesR. ChiPseq_HD: ChIP-seq peaks + HD SNPs. Coding_HD: coding variants + HD SNPs. mQTL_HD: mQTLs + HD SNPs. sQTL_HD: sQTL variants + HD SNPs. Cons100w_HD: conserved variants across 100 vertebrates + HD SNPs. Finemap80k_HD: finely mapped variants + HD SNPs.

### Genomic prediction of traits

In total, we evaluated the genomic prediction accuracy in 216 scenarios, across 6 single-trait analysis, 6 functional categories, 4 breeds in the validation population, and 2 Bayesian methods. Out of these 216 scenarios, 142 (66%) times, HD SNPs combined with functional variants increased genomic prediction accuracy, compared to the prediction only using the HD SNPs (Figure 3 and 4). In 51 out of 216 times (24%), the increase in prediction accuracy ([*r*_*functional*_ − *r*_*HD*_] × 100%) was greater than 1%. These 51 cases were almost all accounted for by Jersey (15/51) and Australian Red (34/51), with only 2 cases in Holstein cattle. In 29 analyses (14%), the increase in prediction accuracy over HD SNPs was greater than 2%. All these 29 cases were for non-Holstein breeds. Amongst tested functional sets, genomic prediction accuracy was the best when the HD variants were combined with conserved variants (Cons100w_HD). In contrast, averaged across tested scenarios, adding randomly selected 944,000 variants to HD had a slightly worse or no improvement in prediction accuracy (−0.5%±0.49%) compared to only using the HD panel to predict.

**Figure 3.**
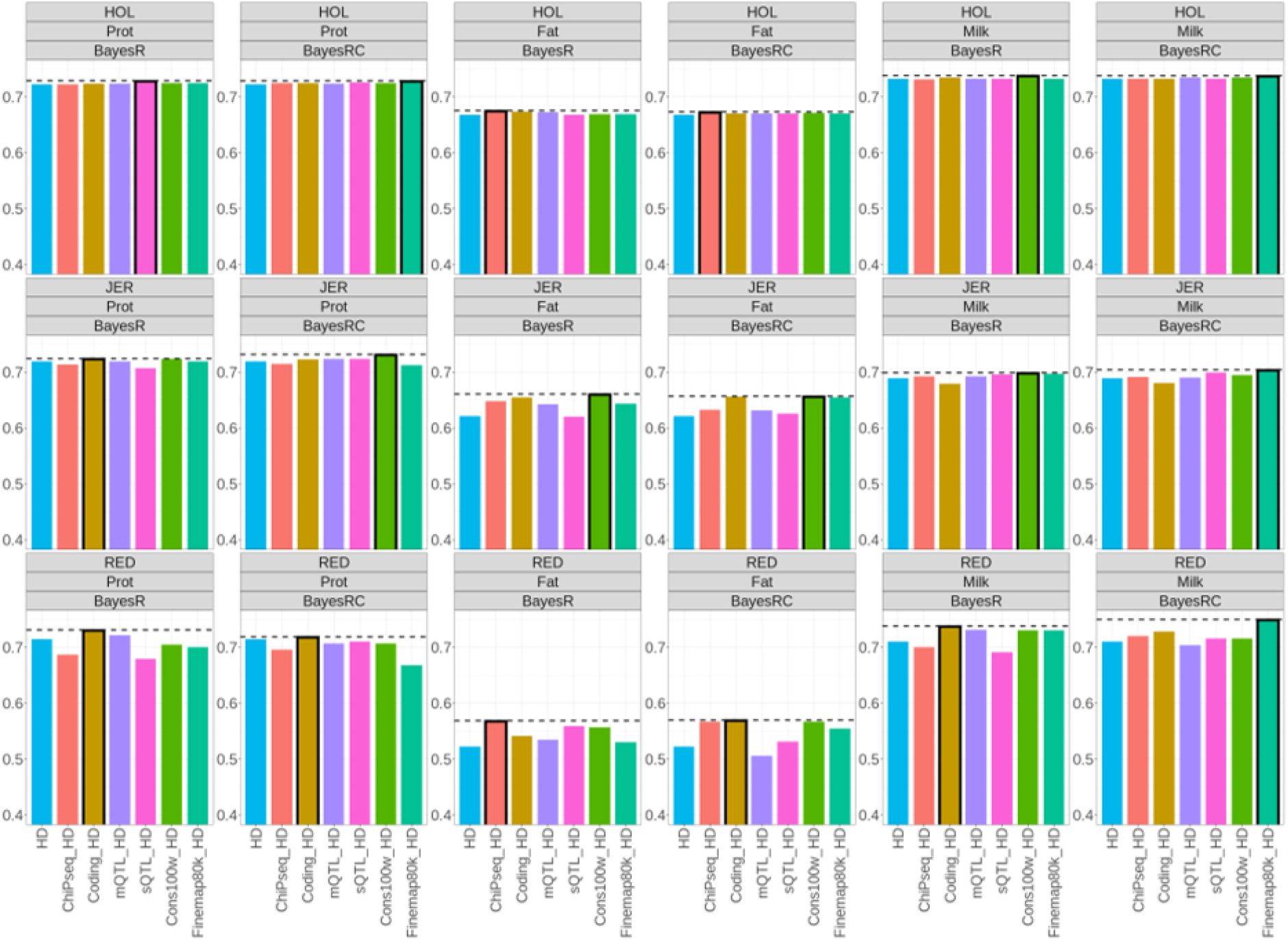
Genomic prediction accuracy (Pearson correlation coefficient, Y-axis) for production traits, across different functional/evolutionary variant sets, breeds and Bayesian methods. A black border and a dashed line of a bar indicate that it has the highest genomic prediction accuracy in the panel. HOL: Holstein breed. JER: Jersey breed. RED: Prot: milk protein yield. Fat: milk fat yield. Milk: milk yield. Australian Red. ChiPseq_HD: ChIP-seq peaks + HD SNPs. Coding_HD: coding variants + HD SNPs. mQTL_HD: mQTLs + HD SNPs. sQTL_HD: sQTL variants + HD SNPs. Cons100w_HD: conserved variants across 100 vertebrates + HD SNPs. Finemap80k_HD: finely mapped variants + HD SNPs.

**Figure 4.**
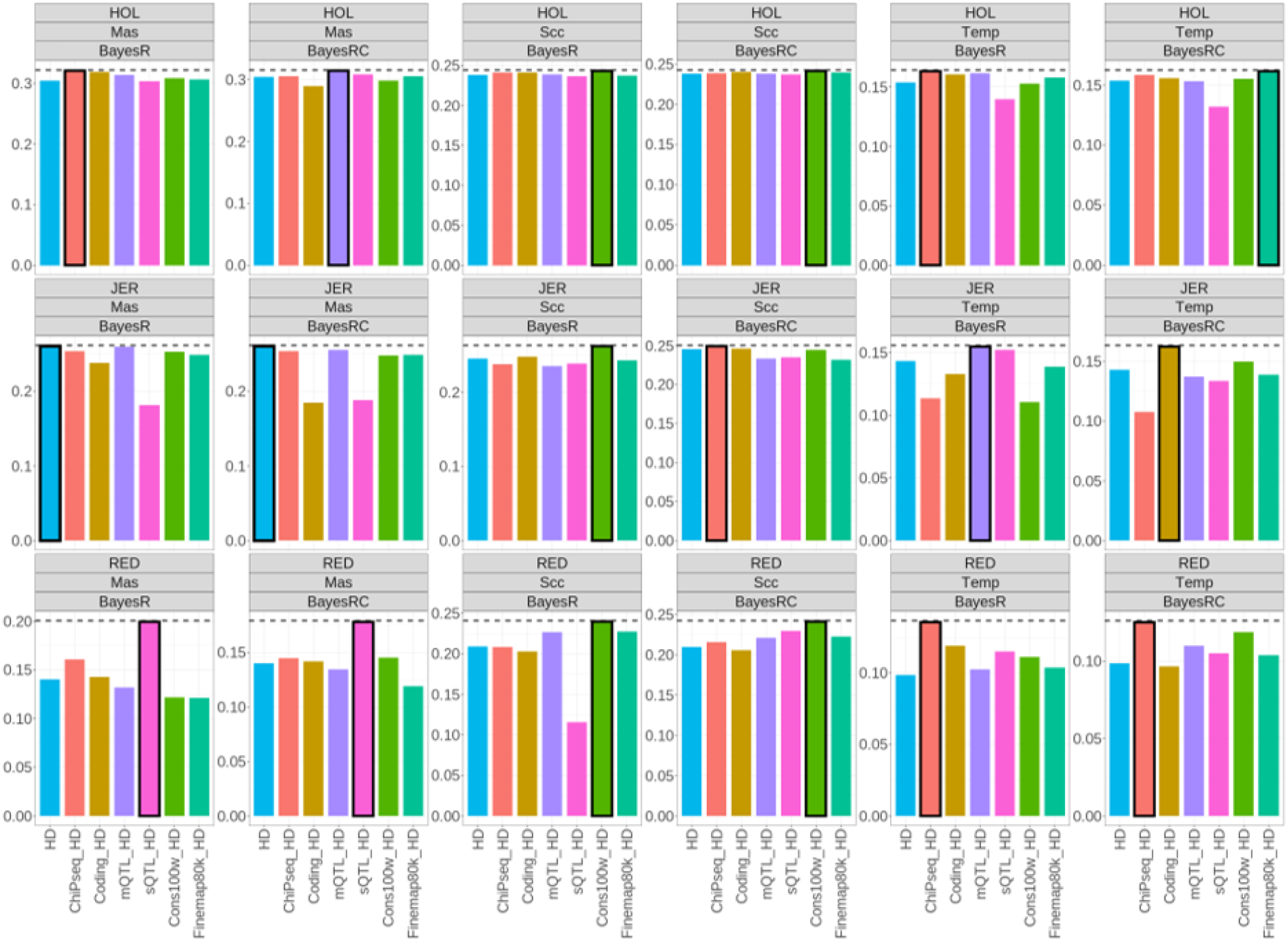
Genomic prediction accuracy (Pearson correlation coefficient, Y-axis) for mastitis, somatic cell count and temperament across different functional/evolutionary variant sets, breeds and Bayesian methods. A black border and a dashed line of a bar indicate that it has the highest genomic prediction accuracy in the panel. HOL: Holstein breed. JER: Jersey breed. RED: Australian Red. Mas: mastitis. Scc: somatic cell count. Temp: temperament. ChiPseq_HD: ChIP-seq peaks + HD SNPs. Coding_HD: coding variants + HD SNPs. mQTL_HD: mQTLs + HD SNPs. sQTL_HD: sQTL variants + HD SNPs. Cons100w_HD: conserved variants across 100 vertebrates + HD SNPs. Finemap80k_HD: finely mapped variants + HD SNPs.

As shown in Figure 3, the genomic prediction accuracy of milk production traits using HD SNPs in Holstein cattle was already high (around 0.7) and the increases in accuracy from functional variants were very small. However, larger increases were evident in Jersey and Australian Red. For milk production traits, 10 out of 18 times the genomic prediction accuracy was the most improved by conserved variants and coding variants combined with HD SNPs, followed by finely mapped variants combined with HD SNPs (4/18), ChIP-seq tagged variants (3/18) combined with HD SNPs. sQTL combined with HD variants had the highest accuracy when predicting protein yield in Holstein.

As shown in Figure 4, the greatest increases in prediction accuracy for traits mastitis, somatic cell count and temperament were again seen in non-Holstein breeds. Chip-seq peak tagged variants combined with HD SNPs (5/18 times) and conserved variants combined with HD SNPs (5/18 times) had the best performances in predicting mastitis, somatic cell count and temperament.

Across all scenarios, we did not see a clear distinction in prediction accuracy between BayesR and BayesRC in the current study. There may be some tendencies where BayesRC had a higher accuracy than BayesR for somatic cell count, mastitis and temperament.

However, none of these differences were significant.

## Discussion

Our systematic evaluations show that functional information can improve genomic mapping and prediction of cattle traits, even when HD SNPs are used, although there were times where HD SNPs alone still had robust performances. It is usually the less represented breeds, such as Jersey and Australian Red who benefited the most from the improvements using functional data. This suggests functional information can well complement HD SNPs especially in breeds with smaller training sets. Adding randomly selected variants to the HD panel reduces mapping precision and provided no improvement in prediction accuracy compared to only using the HD panel. This supports that the The benefit provided via selecting variants based on functional importance can not simply be achieved by adding more sequence variants.

We show that the biological information content which can be used to benefit mapping and/or prediction is different between functional datasets. One of the top-performing functional variant sets in mapping large-effect variants was the finely mapped 80,000 variants (Xiang *et al.*, 2021). This result is somewhat expected as these variants combined information from multiple functional datasets and also included variants affecting multiple dairy cattle traits.

These finely mapped 80,000 variants outperformed the SNPs from the 50K panel in previous evaluations (Xiang *et al.*, 2021). Furthermore, finely mapped 80,000 variants showed enhanced enrichment of large-effect variants and improvement in mapping precision when modelled with BayesRC. This suggests that this much more refined set of variants (chosen because they were more relevant to the traits of interest) are likely more enriched for variants that are more strongly associated with the trait or are causal. BayesRC would only outperform BayesR when there is strong enrichment for QTL in at least one of the defined classes. The other functional groups tested are not trait specific (except mQTL for fat) so likely less enriched relative to each trait.

Previous results showed that coding-related variants did not explain a significant amount of heritability (Koufariotis *et al.*, 2018; Xiang *et al.*, 2019). In the current study, coding-related variants combined with HD SNPs showed enhanced enrichment with large-effect variants and improvement in mapping precision. This implies that variants affecting protein coding may not necessarily be good at capturing all the genetic variance of polygenic traits. The small set of mQTLs, derived from milk fat showed strong enrichment of large-effect variants but did not show improvement in mapping precision over HD SNPs. This set of variants needs future investigations.

Unlike the results in mapping large-effect variants, for genomic prediction, the top-performing variant set is the conserved variants combined with HD SNPs. The advantage of adding conserved variants to HD SNPs was particularly evident when predicting somatic cell count, mastitis and temperament of non-Holstein breeds (Figure 4). In fact, in these scenarios HD SNPs alone did not perform so well and this leaves more room for functional variants to improve the prediction accuracy. Another variant set that performed well in genomic prediction is the set of ChIP-seq peak tagged variants. Again, such an advantage was the most evident when predicting somatic cell count, mastitis and temperament in non-Holstein breeds. Interestingly, ChIP-seq variants combined with HD SNPs appear to show some particular advantages in predicting temperament. There may be some large-effect variants for temperament captured by ChIP-seq peaks.

We found that sQTL variants combined with HD SNPs had variable performances in mapping and prediction. This set did not show good performance in detecting enrichment of informative variants, but overall significantly increased mapping precision over HD SNPs. In genomic prediction, its performance was not impressive. This is somewhat different from previous studies which showed that sQTLs are enriched with complex trait QTL(Li *et al.*, 2016; Xiang *et al.*, 2019; Xiang *et al.*, 2018). One explanation is that sQTLs or any other eQTLs were not trait specific and are plagued by LD, which is particularly strong for Holstein breeds that dominated the discovery population. Another explanation is that the sample size with which we used to discover sQTLs is still small (N~120) and we should re-discover and re-evaluate this set of variants when there is a larger sample size.

As mentioned earlier, BayesRC would only outperform BayesR when there is strong enrichment for QTL in at least one of the defined classes. It would also require functional information to be trait-specific. We saw advantages in BayesRC over BayesR in detecting enrichment with large-effect variants using finely mapped variants, coding variants and mQTLs. BayesRC also had advantages over BayesR in mapping precision when used with finely mapped variants and coding variants. While these functional data are expected to be informative, they did not provide consistent advantages for BayesRC to predict traits over BayesR. Across all tested cases, we did not see strong advantages in BayesRC over BayesR in genomic prediction (Figure 4). BayesRC may have some tendencies to better predict somatic cell count, mastitis and temperament than BayesR. However, the differences were not statistically significant. The reason behind these observations may be complex.

We know that not all variants in the functional datasets are informative and many sequence variants are in strong LD. BayesR and BayesRC both have limitations where variants are in very strong LD. In addition, if most causal variants are quite well tagged by HD variants and if validation animals are highly related to the discovery animals, the room to improve prediction accuracy is limited. Also, there may be less common variants that are not tagged by HD SNPs, but these variants are not well imputed. Further, the optimal tissues and/or experimental conditions to generate functional data that can be better used for improving genomic prediction are usually not known. Therefore, the marriage between functional data and genomic prediction is still at its very early stage.

We therefore suggest two future research directions to improve on the current results. The first is to increase the information content in functional datasets. This can be achieved by either increasing the sample size (biological replicates, tissues and experimental conditions) of functional datasets or by developing better bioinformatic tools to increase the signal-to-noise ratio in functional datasets before they can be processed by genomic prediction models. The second direction is to improve the current genomic prediction models. Because the type and complexity of functional data will keep growing, it will be necessary to develop more sophisticated and flexible methods to better extract information from complex functional data. For example, an extended BayesRC that can model quantitative biological priors, instead of qualitative classes will be needed. Similarly, in the future we will use larger sample sizes and diverse breeds in the training model to reduce LD between sequence variants. This will also increase the need for Bayesian methods to be more efficient.

In conclusion, our evaluation of Bayesian genomic prediction using functional and evolutionary information with HD SNPs provides novel insights into this emerging area. We show that functional datasets of conserved variants, coding variants, ChIP-seq peaks and previously finely mapped variants can improve genomic mapping and/or genomic prediction, even when HD SNPs are used. Such improvements usually benefit non-Holstein breeds, given the current available functional datasets. We found that by using informative biological priors, BayesRC has significant advantages over BayesR in detecting enrichment with large-effect variants and in mapping precision. However, the advantage of BayesRC over BayesR for genomic prediction was not consistent. Our results highlight the need to develop better tools to extract information from complex functional datasets which will benefit genomic prediction in large datasets. Fusing functional genomics with genomic selection presents great opportunities to develop new technologies that improve animal breeding and genetics.

## Acknowledgments

Australian Research Council’s Discovery Projects (DP160101056 and DP200100499) supported R.X. and M.E.G. DairyBio, a joint venture project between Agriculture Victoria (Melbourne, Australia), Dairy Australia (Melbourne, Australia) and the Gardiner Foundation (Melbourne, Australia), funded computing resources used in the analysis. The authors also thank the University of Melbourne, Australia for supporting this research. No funding bodies participated in the design of the study nor analysis, or interpretation of data nor in writing the manuscript. DataGene and CRV provided access to the reference data used in this study. We thank Gert Nieuwhof, Kon Konstantinov and Timothy P. Hancock (DataGene) and staff from DairyNZ for the preparation and provision of data. We thank Dr. Sunduimijid Bolormaa for the sequence variant data imputation. We thank Drs. Iona M. MacLeod and Hans D. Daetwyler for critical reading of the manuscript.

